# EXPLORING THE DISTRIBUTION OF SINGLE NUCLEOTIDE POLYMORPHISMS ACROSS HUMAN EXONS AND INTRONS

**DOI:** 10.1101/2024.03.23.586436

**Authors:** Magdalena Fraszczak, Jakub Liu, Magda Mielczarek, Paula Dobosz, Joanna Szyda

## Abstract

Among all types of mutations, single nucleotide polymorphisms are the most common type of genomic variation. In our study, we explore the counts of single nucleotide polymorphisms in particular exons and introns of the human genome based on the data set of 1,222 individuals of Polish origin that comprises 41,836,187 polymorphisms. In particular, chromosomes 1 and 22 were considered as representatives of two markedly different DNA molecules, since HSA01 represents the longest and HSA22 is one of the shortest chromosomes. The results demonstrate that outer (first, last) exons as well as the first introns harbour significantly more SNPs than other genic regions. The observed differences in counts reflect the distinct functional roles of those genomic units.

## INTRODUCTION

Single nucleotide polymorphisms (SNPs) are the most common type of genomic variation not only in humans but also in several other species. Still, their genomic distribution is not random (1, 2). Although they are located in all functional genomic elements (promoters, exons, introns, 5’ and 3’ UTR, intergenic regions) their density varies between regions, with exons and splice sites (defined by exon-intron boundaries) being the most conservative, that is, SNP-sparse (3). But even within functional genomic units with a sequential structure, such as introns and exons, the density of SNPs is highly non-uniform (4), with clusters of adjacent SNPs being an often-observed characteristic of the human genome. In particular, Hodgkinson and Eyre-Walker (5) and Prendergast et al. (6) estimated an excess of intronic SNPs located in a single-bp proximity to each other Matsushita and Kano-Sueoka (7) recently reported differential clustering of synonymous and non-synonymous SNPs among consecutive exons of the human *HLA-A* gene, while in the whole genome scope Back and Walther (8) observed in *Arabidopsis thaliana* a higher density of SNPs located in the first intron than in the next introns. It has been widely agreed in the literature that such non-random distribution of SNPs must have evolutionary impactions and is the result of mutational hotspots, or that SNP clusters arise due to structural properties of DNA that mechanically promote the accumulation of such point mutations.

It is important to note that SNP density and SNP count shall be regarded as two non-equivalent measures of SNP genomic distribution. The SNP count expresses a raw number of polymorphisms identified within a functional unit regardless of the unit length and distances between SNPs, while the SNP density is represented by various descriptive statistics (such as the mean or median) of pairwise distances between adjacent SNPs and is only indirectly related to the number of polymorphisms. In our analysis, we explored the number of SNPs in the human genome, focusing on differences in SNP counts among consecutive introns and exons of a given gene. The underlying hypothesis that we aim to verify is that there are differences in the numbers of SNPs among particular exons and introns that may reflect the differential role of particular exons and introns in the formation of the final product of a gene (mRNA). For this purpose, we used a large data set of whole genome sequences of 1,222 persons representing the 1000 Polish Genomes database (9).

## MATERIAL AND METHODS

### Population studied

The cohort analysed consisted of 41,836,187 SNPs identified in genomes of 1,222 individuals of Polish origin, recruited during the “Search for Genomic Markers Predicting the Severity of the Response to COVID-19” project. The sample consisted of 697 men and 525 women of age by sampling between 2 and 99 years old, with a mean age of 45 years. All samples were collected between April 2020 and April 2021. Details on subject ascertainment, whole genome sequencing, and variant calling were described by (9).

### Variant filtering and genomic annotation

Filtering of the initial set of 41,836,187 SNPs was performed using Vcftools software (10) that was used to compose the subsets of SNPs located in HSA01 and HSA22. These two chromosomes were selected for downstream analysis as representatives of two markedly different DNA molecules in the human, since HSA01 represents the longest and HSA22 is one of the shortest chromosomes. In particular, HSA01 consists of 248.96 Mbp, which represents almost 8.04% of the genome and contains 5,485 genes. In contrast, HSA22 consists of only 50.82 Mbp, representing 1.64% of the genome, and contains 1,258 genes (GRCh38.p14 assembly GCA_000001405.29). Furthermore, variants with a mapping quality score of at least 20 were discarded. The remaining variants were genomically annotated using the Ensembl Variant Effect Predictor tool (11). The final selection of SNPs that were subjected to downstream analysis consisted of variants located in introns or exons of canonical transcripts of each gene.

### Data exploration

The statistical analysis pipeline was set up to follow the hypothesis testing scheme of increasing biological specificity, that was applied separately for exons and introns, as well as separately for each group of genes defined by the same numbers of exons/introns. At each testing step, the null-hypothesis was rejected based on the nominal type I error rate ≤ 0.05.

1. The null-hypothesis of the total number of SNPs being equal among genes was tested using the 2 goodness of fit test calculated using the *chisq*.*test* function implemented in the R *stats* library.

2. The null-hypothesis of the number of SNPs being equal in each exon/intron was tested using the Friedman test (12) implemented using the *PMCMRplus* package in R.

3. For the groups of genes with significant differences in SNP numbers tested in step 2), the null-hypothesis of no differences in SNP numbers between each possible pairs of exons/introns was tested using the Conover test (13) implemented using the *PMCMRplus* package in R.

## RESULTS

### Genomic distribution of SNPs on HSA01 and HSA22

Among the 41,836,187 SNPs identified in our data set of 1,222 persons, 1,705,575 SNPs were located in genes (i.e. exons or introns) on HSA01 and HSA22, which made up 4.06% of all SNPs. 5,177 of genes on HSA01 and 1,313 of genes on HSA22 contained at least one SNP, what resulted in the number of SNPs per gene ranging from 1 to 13,124 (HSA01) and from 1 to 8,083 (HSA22). The average number of SNPs per gene amounted to 303±779 on HSA01 and 225±571 on HSA22, with AGBL4 (ENSG00000186094) being the most SNP-rich gene on HSA01 that contained 13,124 SNPs, most of which was located in the third intron. Most of the genes, that is, 1,681 on HSA01 and 461 on HSA22 contained only one exon (Supplementary Figure 1). In all downstream analyses, to maintain class counts that allow a reasonable estimation of type I and type II errors, genes with 3 to 12 exons were considered for further analysis. As expected biologically, in both considered chromosomes, exons contained fewer SNPs than introns, the average number of SNPs per exon was the highest for genes with a low number of exons, while the average number of SNPs per intron, that varied between 20 and 90 SNPs, did not depend on the number of introns in a gene (Figure1). 1,074 of those SNPs represented the HIGH impact class as defined by the Sequence Ontology. On HSA01, 507 of them were located in exons and 370 in introns, but a reverse pattern was observed on HSA22 with 97 high-impact SNPs in exons and 104 SNPs in introns.

**Figure 1.**
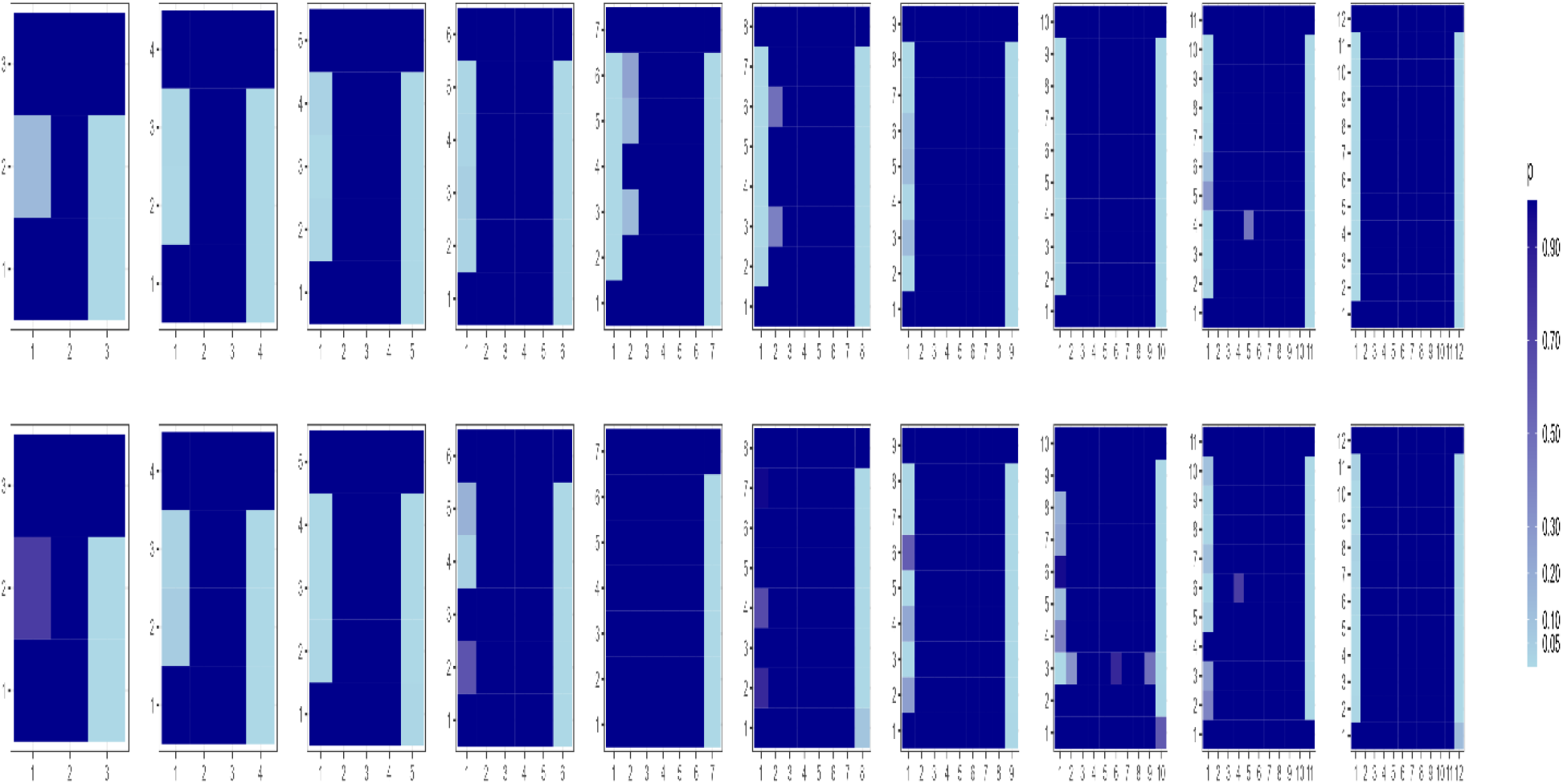
The average number of SNPs on HSA01 and HSA22 located in exons or introns.

### Differences in the number of SNPs located in exons

Within all gene groups, defined by their exon counts, the total numbers of SNPs in exons were highly significantly different. Also, the differences in SNP count tested between exons were highly significant for all gene groups and both chromosomes (Supplementary Table 1). Pairwise differences in SNP counts between exons were visualised in Figure 2. For HSA01, a very consistent pattern emerged, showing the significant excess of SNPs in the first and the last exon, but the first exons always contained fewer SNPs than in the last. The same pattern was also true for HSA22, except genes represented by 7 and 8 exons. Considering only the subset of HIGH impact SNPs (defined by Sequence Ontology), 29% of them (HSA01) and 35% (HSA22) were assigned to the last exon.

**Figure 2.**
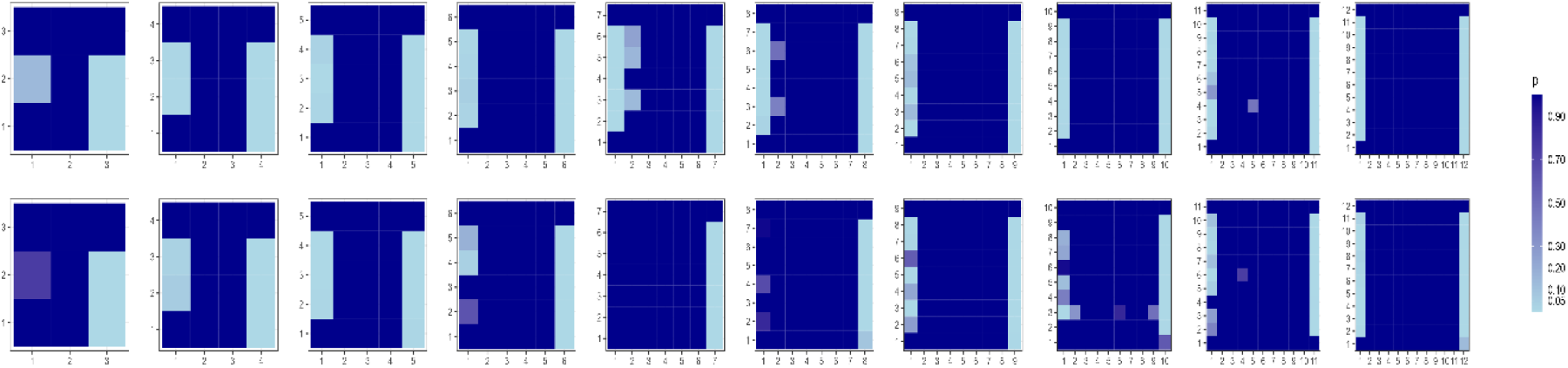
Visual representation of the significance of pairwise comparisons of SNP counts in particular exons. P values correspond to testing the alternative hypothesis of a lower number of SNPs within the i-th exon represented by the Y-axis (rows) than within the j-th exon represented by the X-axis (columns). Top HSA01, bottom HSA22.

### Differences in the number of SNPs located in introns

Within all gene groups, the total numbers of SNPs in introns were highly significantly different. The differences in SNP count tested between introns were highly significant for all gene groups on HSA01, while on HSA22 there were no significant differences between introns for genes representing groups of 3, 5, and 6 introns (Supplementary Table 1). Pairwise comparisons between introns yielded a very consistent pattern for HSA01, showing significantly higher numbers of SNPs harboured by the first intron. On HSA22 this pattern was also observed, albeit being somewhat less consistent across all possible pairwise comparisons (Figure 3). 26% and 27%of HIGH impact SNPs on HSA01 and HSA22 respectively were assigned to the 1st intron, while 27% (HSA01) and 31% (HSA22) were located in the last intron.

**Figure 3.**
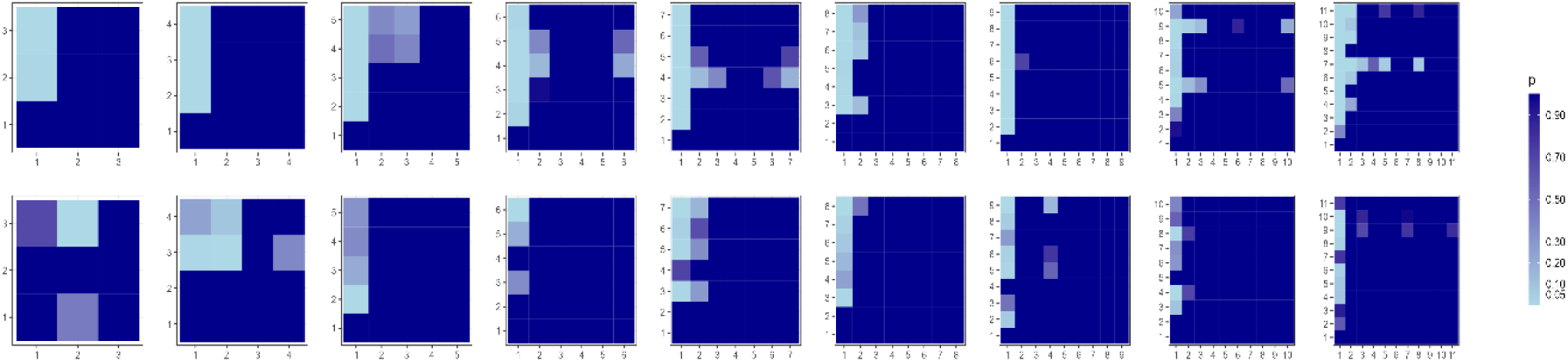
Visual representation of the significance of pairwise comparisons of SNP counts in particular introns. P values correspond to testing the alternative hypothesis of a lower number of SNPs within the i-th exon represented by the Y-axis (rows) than within the j-th exon represented by the X-axis (columns). Top HSA01, bottom HSA22.

## DISCUSSION

Today, very large data sets of SNPs identified from whole genome sequencing are available, such as e.g. the huge resource provided by the UK Biobank (ukbiobank.ac.uk). Still, the data set analysed in our study possesses characteristics that make it advantageous for the SNP count analysis. In particular, all 1,222 individuals were ascertained and processed as a single cohort and therefore underwent identical methodology of variant calling, including the genotyping platform and sequence pre-processing, which allowed for minimising the technical bias of SNP calling. Moreover, this data set represents a timely and geographically uniform group of individuals of Polish origin (9) and therefore excludes the ascertainment bias of SNP frequency due to population stratification and selection.

### SNP distribution

In our study, we deliberately focused on SNP count instead of SNP density, even though the length of particular exons and introns varies considerably. However, while some studies have suggested a correlation between gene length or exon/intron count and SNP density (14), the relationship is not always straightforward. Gene function, selective pressures, and genomic context can influence SNP counts (3). Therefore, introns and exons were in our study regarded as functional genomic units, and not as a mechanical sequence of nucleotides.

The functional role of the genomic region strongly determines the localization of polymorphisms, since SNPs in exons have a potential impact on gene products that on a further scope may be the cause of a disease or may alter quantitative phenotypes (15). Since introns exhibit various regulatory roles, the presence of polymorphism in introns may indirectly impact gene products or their expression levels (16). Still, due to the generally more severe potential consequences of polymorphisms in exons, the expectation is that exons contain fewer SNPs than introns (17), which was confirmed by our study. Moreover, genes with a low number of exons had the highest mean number of SNPs per exon, what was also observed in this study. This may be related to the fact that smaller genes are frequently expressed during an individual’s lifetime because they are typically involved in functions that require fast responses, such as the immune system. These specific functions contribute to a higher variation which facilitates the response to and interaction with changing environment (14).

### SNP counts in introns

The biological role of introns is manyfold. They allow for alternative splicing (18) but additionally influence the stability of mRNA molecules (19) and themselves contain noncoding RNA genes (20) and regulatory elements, especially enhancers that affect the rate of transcription, known as the phenomenon of Intron Mediated Enhancement (21, 22). In our study, the significant excess of SNPs was observed in first introns. Of all the introns, the first one has been recognized as having special features and functions including, among others, correcting cytoplasmic localization of some mRNAs as well as transcriptional and translational regulation (23). The important role of genetic variation in the first introns can also be anticipated by observing a very high number (over 120 since 2001) of publications reporting associations of SNPs located in the first intron with a variety of phenotypes measured in humans, animals, and plants (based on PubMed access on 10.01.2024). These specific roles may explain why the human DNA’s first intronic sequence is considered the longest and highly dense of regulatory chromatin marks (24, 25) Considering the criterion of SNP density, the above-mentioned studies identified the first introns as the most conserved regions, that however, provided a highly non-uniform distribution of SNPs along the intronic sequence (26) does not directly translate to the total low number of SNPs. Interestingly, in *Arabidopsis thaliana*, Back and Walther (8) observed that the first introns harbour more SNPs than the subsequent introns.

### SNP counts in exons

In the aforementioned study of Back and Walther (8) that used *Arabidopsis thaliana* as the model genome, a high positive correlation was estimated between the sequence variation in the first exons and gene expression, which would explain higher genomic variability of the first exon observed in our study. Also, in a slightly different context of across-species comparison based on the reference genome sequence, Castle (4) observed higher variability of coding regions in the proximity of the start and stop codons, so the ones typically corresponding to the first and the last exons, that is in line with our observation of SNP excess in the first and the last exons. Moreover, first and last exons include not only the protein-coding sequence but also the 5′- and 3′-untranslated regions (UTRs). The 5′UTR is the RNA sequence immediately upstream of the coding RNA. It is generally not a translated region but, due to the importance of the sequence for RNA transcription, stability, and translation, genetic variants modifying these elements are likely to have a profound effect. Analogously, the 3′UTR is located downstream of the coding sequence, and it is involved in regulatory processes, including RNA stability, mRNA translation, and localization. The 3′ UTR is characterized by binding sites for microRNAs and RNA-binding proteins and thus any variation of this region may lead to a change in gene expression (27). Although, UTRs are considered to have approximately the same genomic footprint as protein-coding regions still, polymorphisms within coding sequences may directly affect proteins and thus affect their function. This explains that higher SNP number in the first and last exons may be related to the presence of UTRs.

## Conclusions

The distribution of single nucleotide polymorphisms among introns and exons is not only highly non uniform, but also exhibits a very consistent pattern of first introns, first exons and last exons harbouring significantly more polymorphisms. This observation reflects the distinct functional role of those genomic units.

## DATA AVAILABILITY

Summary statistics of single nucleotide polymorphisms characterized for the whole genomes of the individuals were provided by Kaja et al. (9) on https://github.com/MNMdiagnostics/NaszeGenomy.

## SUPPLEMENTARY DATA

Supplementary Data are available at NAR online.

## AUTHOR CONTRIBUTIONS

Magdalena Fraszczak: Formal Analysis, Methodology, Visualization. Jakub Liu: Data curation, Formal Analysis, Software. Magda Mielczarek: Supervision, Writing. Paula Dobosz: Funding acquisition, Resources, Writing. Joanna Szyda: Conceptualization, Methodology, Writing.

## ACKNOWLEDGEMENTS

The computational power was provided by Poznan Supercomputing and Networking Centre. The Authors would like to thank all sample donors that participated in the study, as well as the medical personnel of the Central Clinical Hospital of the Ministry of the Interior and Administration in Warsaw for their active support. The idea for this study was raised in August 2022, bioinformatics analysis performed in autumn 2022 and the manuscript written in 2023. The datasets presented in this study can be found in an online repository: https://1000polishgenomes.com [access date: October 2022]. Full cohort description can be found in the following paper: https://doi.org/10.3390/ijms23094532.

## REFERENCES

1. Neininger, K., Marschall, T. and Helms, V. (2019) SNP and indel frequencies at transcription start sites and at canonical and alternative translation initiation sites in the human genome. PLoS One, 14, e0214816.

2. Amos, W. (2010) Even small SNP clusters are non-randomly distributed: is this evidence of mutational non-independence? Proc. R. Soc. B Biol. Sci., 277, 1443–1449.

3. Deng, N., Zhou, H., Fan, H. and Yuan, Y. (2017) Single nucleotide polymorphisms and cancer susceptibility. Oncotarget, 8, 110635–110649.

4. Castle, J.C. (2011) SNPs Occur in Regions with Less Genomic Sequence Conservation. PLoS One, 6, e20660.

5. Hodgkinson, A. and Eyre-Walker, A. (2010) Human Triallelic Sites: Evidence for a New Mutational Mechanism? Genetics, 184, 233–241.

6. Prendergast, J.G.D., Pugh, C., Harris, S.E., Hume, D.A., Deary, I.J. and Beveridge, A. (2019) Linked Mutations at Adjacent Nucleotides Have Shaped Human Population Differentiation and Protein Evolution. Genome Biol. Evol., 11, 759–775.

7. Matsushita, T. and Kano-Sueoka, T. (2023) Non-random Codon Usage of Synonymous and Non-synonymous Mutations in the Human HLA-A Gene. J. Mol. Evol., 91, 169–191.

8. Back, G. and Walther, D. (2021) Identification of cis-regulatory motifs in first introns and the prediction of intron-mediated enhancement of gene expression in Arabidopsis thaliana. BMC Genomics, 22, 390.

9. Kaja, E., Lejman, A., Sielski, D., Sypniewski, M., Gambin, T., Dawidziuk, M., Suchocki, T., Golik, P., Wojtaszewska, M., Mroczek, M., et al. (2022) The Thousand Polish Genomes—A Database of Polish Variant Allele Frequencies. Int. J. Mol. Sci., 23.

10. Danecek, P., Auton, A., Abecasis, G., Albers, C.A., Banks, E., DePristo, M.A., Handsaker, R.E., Lunter, G., Marth, G.T., Sherry, S.T., et al. (2011) The variant call format and VCFtools. Bioinformatics, 27, 2156–2158.

11. McLaren, W., Gil, L., Hunt, S.E., Riat, H.S., Ritchie, G.R.S., Thormann, A., Flicek, P. and Cunningham, F. (2016) The Ensembl Variant Effect Predictor. Genome Biol., 17, 122.

12. Friedman, M. (1937) The Use of Ranks to Avoid the Assumption of Normality Implicit in the Analysis of Variance. J. Am. Stat. Assoc., 32, 675–701.

13. Conover, W J, and Iman, R.L. (1979) Multiple-comparisons procedures. Informal report. United States:

14. Lopes, I., Altab, G., Raina, P. and de Magalhães, J.P. (2021) Gene Size Matters: An Analysis of Gene Length in the Human Genome. Front. Genet., 12, 559998.

15. Nair, V., Sankaranarayanan, R. and Vasavada, A.R. (2021) Deciphering the association of intronic single nucleotide polymorphisms of crystallin gene family with congenital cataract. Indian J. Ophthalmol., 69, 2064–2070.

16. Mukherjee, D., Saha, D., Acharya, D., Mukherjee, A., Chakraborty, S. and Ghosh, T.C. (2018) The role of introns in the conservation of the metabolic genes of Arabidopsis thaliana. Genomics, 110, 310–317.

17. Frigola, J., Sabarinathan, R., Mularoni, L., Muiños, F., Gonzalez-Perez, A. and López-Bigas, N. (2017) Reduced mutation rate in exons due to differential mismatch repair. Nat. Genet., 49, 1684–1692.

18. Bush, S.J., Chen, L., Tovar-Corona, J.M. and Urrutia, A.O. (2017) Alternative splicing and the evolution of phenotypic novelty. Philos. Trans. R. Soc. B Biol. Sci., 372, 20150474.

19. Gupta, S.K., Carmi, S., Ben-Asher, H.W., Tkacz, I.D., Naboishchikov, I. and Michaeli, S. (2013) Basal Splicing Factors Regulate the Stability of Mature mRNAs in Trypanosomes . J. Biol. Chem., 288, 4991–5006.

20. Chorev, M. and Carmel, L. (2012) The function of introns. Front. Genet., 3, 55.

21. Clancy, M. and Hannah, L.C. (2002) Splicing of the Maize Sh1 First Intron Is Essential for Enhancement of Gene Expression, and a T-Rich Motif Increases Expression without Affecting Splicing. Plant Physiol., 130, 918–929.

22. David-Assael, O., Berezin, I., Shoshani-Knaani, N., Saul, H., Mizrachy-Dagri, T., Chen, J., Brook, E. and Shaul, O. (2006) AtMHX is an auxin and ABA-regulated transporter whose expression pattern suggests a role in metal homeostasis in tissues with photosynthetic potential. Funct. Plant Biol., 33, 661–672.

23. Jo, B.-S. and Choi, S.S. (2015) Introns: The Functional Benefits of Introns in Genomes. Genomics Inf., 13, 112–118.

24. Park, S.G., Hannenhalli, S. and Choi, S.S. (2014) Conservation in first introns is positively associated with the number of exons within genes and the presence of regulatory epigenetic signals. BMC Genomics, 15, 526.

25. Jo, S.-S. and Choi, S.S. (2019) Analysis of the Functional Relevance of Epigenetic Chromatin Marks in the First Intron Associated with Specific Gene Expression Patterns. Genome Biol. Evol., 11, 786–797.

26. Majewski, J. and Ott, J. (2002) Distribution and characterization of regulatory elements in the human genome. Genome Res., 12, 1827–1836.

27. Steri, M., Idda, M.L., Whalen, M.B. and Orrù, V. (2018) Genetic variants in mRNA untranslated regions. Wiley Interdiscip. Rev. RNA, 9, e1474.

