## Supplementary data 1 for "EXPLORING THE DISTRIBUTION OF SINGLE NUCLEOTIDE POLYMORPHISMS ACROSS HUMAN EXONS AND INTRONS"

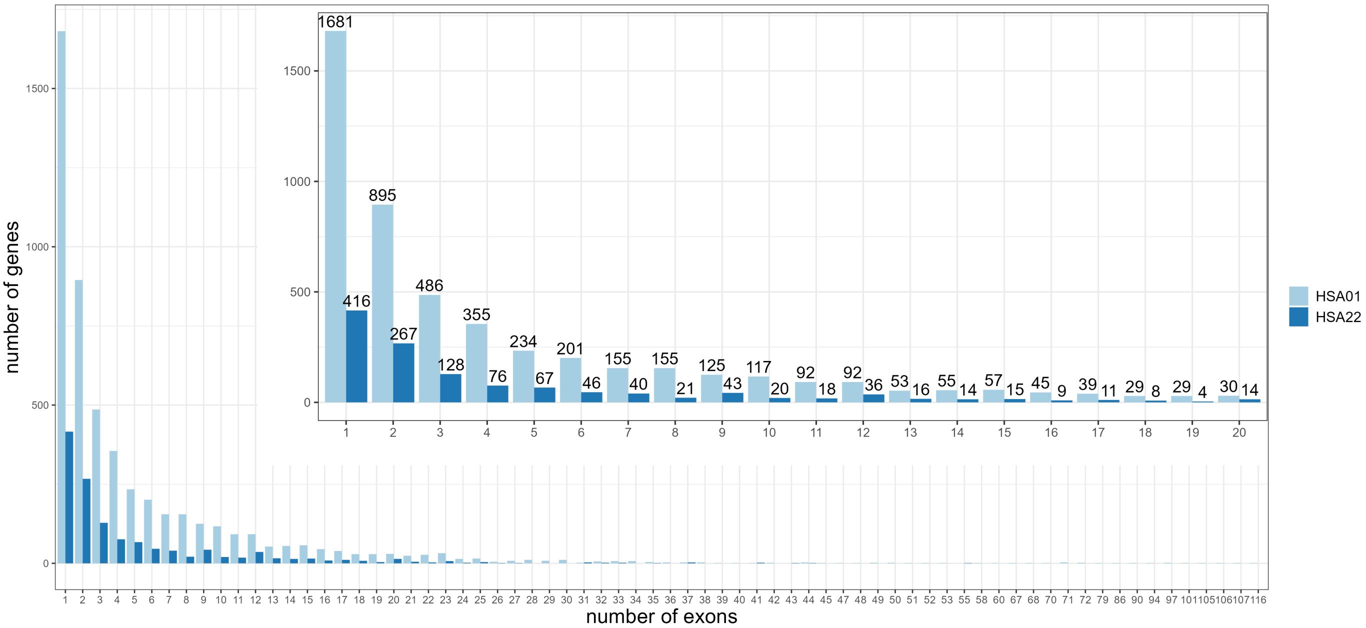

Figure 1. Counts of genes on HSA01 and HSA22 with the given number numbers of exons.

| no exons | exons HSA1 | exons HSA22 | introns HSA1 | introns HSA22 |
|---|---|---|---|---|
| 3 | $9.77 \cdot 10^{-59}$ | $1.11 \cdot 10^{-18}$ | - | - |
| 4 | $6.59 \cdot 10^{-61}$ | $1.61 \cdot 10^{-11}$ | $8.27 \cdot 10^{-3}$ | $2.56 \cdot 10^{-1}$ |
| 5 | $3.74 \cdot 10^{-52}$ | $3.23 \cdot 10^{-15}$ | $2.49 \cdot 10^{-5}$ | $4.71 \cdot 10^{-4}$ |
| 6 | $3.59 \cdot 10^{-56}$ | $1.87 \cdot 10^{-14}$ | $5.41 \cdot 10^{-9}$ | $1.87 \cdot 10^{-1}$ |
| 7 | $1.74 \cdot 10^{-54}$ | $4.11 \cdot 10^{-9}$ | $3.75 \cdot 10^{-7}$ | $1.93 \cdot 10^{-1}$ |
| 8 | $4.74 \cdot 10^{-52}$ | $5.26 \cdot 10^{-6}$ | $5.57 \cdot 10^{-7}$ | $1.93 \cdot 10^{-1}$ |
| 9 | $9.11 \cdot 10^{-46}$ | $4.64 \cdot 10^{-17}$ | $5.07 \cdot 10^{-6}$ | $1.16 \cdot 10^{-3}$ |
| 10 | $5.35 \cdot 10^{-46}$ | $3.88 \cdot 10^{-9}$ | $9.94 \cdot 10^{-11}$ | $2.79 \cdot 10^{-3}$ |
| 11 | $2.97 \cdot 10^{-40}$ | $4.38 \cdot 10^{-10}$ | $1.28 \cdot 10^{-5}$ | $2.17 \cdot 10^{-2}$ |
| 12 | $6.88 \cdot 10^{-35}$ | $7.11 \cdot 10^{-21}$ | $3.34 \cdot 10^{-14}$ | $1.01 \cdot 10^{-3}$ |

Table 1. The P- values in Friedmann tests.
